# Effect of growth forms of bamboo on the mycorrhizal and fluorescent pseudomonas population

**DOI:** 10.1101/2021.12.08.471877

**Authors:** Solomon Das, Y. P. Singh, Yogesh K. Negi, P. C. Shrivastav

**Author notes:** (Solomon Das); (Y.P. Singh); (Yogesh K. Negi) (P.C. Shrivastav).

## Abstract

Bamboo, also called as the poor man’s timber, is one of the fastest growing giant grass species. Having shallow root system and fast growth rate, the dependability of the plant on the rhizospheric microbial web cannot be denied. The study was conducted to explore the population and seasonal variations of indigenous mycorrhizal fungi as well as the functional diversity of plant growth promoting bacteria, especially fluorescent pseudomonas from the different growth forms of *Dendrocalamus strictus*, the most commonly present bamboo species in Indian sub-continent. In a past study, it was established that the growth forms of *D*. *strictus* which developed over time in their respective locations, were genetically varied. The present research further explores the variations in their respective rhizospheric microbes and looks for the role of plant selection phenomenon. Considerable variation in mycorrhizal structures and in the functional diversity of fluorescent pseudomonas was registered. It seems probable that selection pressure of nutrient deficient condition has created a condition that promoted occurrence of high numbers of P solubilisers which, in turn, boosted the mycorrhizal as well as bamboo growth.

## 1. Introduction

Bamboos, considered as the ‘poor man’s timber, are a means to reclaim degraded land, conserve soil, improve environment and carry out drought proofing. Bamboo plantations are important in Greening India Programme (2010–2020) (NMBTTD Report, 2003; INBAR, 2018). Of all the commonly occurring genera of bamboos, the genus *Bambusa* is the widely distributed in India followed by *Dendrocalamus*. Deogun (1937) has differentiated three major growth forms of *D. strictus* based on edaphic conditions, isolation aspect, temperature and humidity. These growth forms growing in different geographical regions show variations in their appearances, arrangement of clumps and cross-sectional thicknesses of culms. These variations, evolved due to the environmental and other factors, are a matter of investigation. A study conducted by Das et al. (2017) explored the genetic variability among and within growth forms of bamboo using RAPD.

One of the important problems in tropical biomass production is the low-soil-phosphate (P) availability. The consumption of chemical fertilisers has significantly increased in last three decades. A large portion of soluble inorganic phosphate applied to soil as chemical fertiliser is rapidly fixed soon after application and becomes unavailable to plants (Santana et al., 2016). Hence, a sustainable fertilisers management becomes essential roping in the application of those microbial inoculants in the soil that are helpful in making P available to plants. Rhizosphere of bamboo holds the seat of several microbial communities which are beneficial in nature. Among the useful microorganisms, plant growth promoting rhizobacteria (PGPR) and arbuscular mycorrhizal (AM) are important ones and the indispensable part of rhizosphere biota, that when grown in association with the host plants, can stimulate the growth of the host (Thangavelu and Udaiyan, 2006).

*Pseudomonas* are one of the well-known solubilisers of phosphate (Chakraborty et al., 1990), producer of plant growth hormones like IAA, and have broad spectrum antagonistic activity against plant pathogens such as antibiosis (Didick et al., 2000), nutrition or site competition (Ahmad et al., 2005) and siderophore production (Fankem et al., 2006). Some species of fluorescent pseudomonas produce higher levels of hydrogen cyanide (HCN) that is toxic to certain pathogenic fungi (Gloria and Hoadley, 1976). The recent study shows that these species are playing major role in regulating the availability of phosphates to the plants with help of HCN (Rijavec and Lapanje, 2016). These characteristics make bacterial species good candidate for use as seed or root inoculants for biological control of soil borne plant pathogens and to avail poorly available nutrients like P. *D. strictus*, one of the most common bamboo species present in India, is totally unexplored in terms of understanding the status and role of beneficial microbes of its rhizosphere and to tap them for diverse benefits.

Some studies have shown the synergistic interactions between phosphate solubilising bacteria and AM fungus which is a phosphate absorber (Nacoon et al., 2020). These interactions in relation to the growth and macroproliferation of *D. strictus* are not traceable in literature. Such information may be vital as *D. strictus* grows in different agro-climatic zones, especially P deficient soils of India. Use of these two microorganisms together may help in tackling the nutritional limitation in P-deficient tropical soils, since both phosphorus solubilisation and absorption can be operationalised at a time and in the same niche. Generating the information on synergistic effects of mixed inoculation on bamboo growth and P uptake is needed. The compatibility between *P. fluorescens* and AM seems to have a certain degree of specificity; therefore, investigation on most effective combinations is required.

## 2. Material and Methods

### 2.1. Sampling locations

The study was conducted at three locations: first, reserve forest of the FRI, Dehradun (latitudes of 30° 20’N and longitudes of 78° 00’E); second, Shivpuri near Byasi Rishikesh (latitudes of 30° 06’N and longitude of 78° 25’E); and third, Chiriapur range, Haridwar district (between the latitude of 29° 52’N and longitude of 78° 11’E). These locations were selected based on the presence of different growth forms of *D. strictus* (Das et al., 2017). Climatic and soil physico-chemical parameters were also recorded (Table 1).

**Table 1.** Climatic conditions and physico-chemical properties of soils of different locations

| Parameter | FRI campus | Shivpuri | Chiriapur |
| --- | --- | --- | --- |
| Annual rainfall (mm) | 2,060.0 | 1,706.8 | 1,261.5 |
| Min. and max. temperature (°C) | 3.9–33.2 | 0.2–32.1 | 7.5–36.9 |
| Above sea level (m) | 435 | 372 | 314 |
| Clay (%) | 13.4 | 16.2 | 10.6 |
| Silt (%) | 15.4 | 16.1 | 10.4 |
| Sand (%) | 71.2 | 67.7 | 79.0 |
| Textural class | Loam | Loam | Sandy loam |
| Soil reaction (pH) | 6.8 | 6.6 | 6.2 |
| Electrical conductivity (dsm<sup>-1</sup>) | 0.24 | 0.25 | 0.18 |
| Organic carbon (%) | 2.4 | 2.0 | 1.0 |
| Nitrogen (kg/ha) | 375.0 | 204.0 | 168.0 |
| Available P (kg/ha) | 69.9 | 53.8 | 12.5 |
| Available K (kg/ha) | 645.1 | 329.3 | 309.1 |

### 2.2. Microbial Status in the Root Zone of Different Growth Forms of *D. strictus*

#### 2.2.1. Arbuscular Mycorrhizae

The sampling for the indigenous AM fungi was done as adopted for collecting the soil samples. The composite soil sample of each clump (500g) was reduced to about 25ml for estimation of mycorrhizal spores (Wet sieving and decanting method of Gerdemann and Nicholson, 1963; Sugar gradient centrifugation method of Daniel’s and Skipper, 1982) and 250 mg of roots for mycorrhizal colonisation ( staining of roots method of Phillips and Hayman, 1970; measurement of mycorrhizal root colonisation using Grid line - Intersect method of Giovanetti and Mosse, 1980). Spores were identified at generic level using the key of Schenck and Perez (1990).

#### 2.2.2. Fluorescent Pseudomonads

##### 2.2.2.1. Isolation

1. Serial dilutions from 10^−1^ to 10^−6^ were prepared from each soil sample (1g).
2. Samples from dilution 10^−3^ to 10^−5^ were inoculated on King’s B medium using spread plate method.
3. Plates were incubated at 30°C for 24–48 hr.
4. Colonies with yellow green fluorescence identified under UV lamp were picked up by inoculation loop and streaked on fresh King’s B medium plate.
5. Pure isolates were further sub-cultured and maintained on slants (kept in refrigerator for 5°C).

### 2.3. Functional Assessment of Growth and Biocontrol Properties

The growth promoting characteristics of fluorescent pseudomonas such a phosphate solubilizing activity (Pikovskaya, 1948) and Indole Acetic Acid (IAA) production (Bric *et al.*, 1991), and biocontrol activities like siderophore production (Meyer and Abdullah, 1978), HCN production (Millar and Higgins, 1970), and chitinase production (Renwick *et al.*, 1991) were estimated.

### 2.4. Statistical analysis

The numbers of isolates recovered from the different growth forms assigned to each class of growth (phosphate solubilising efficiency and IAA production) and bio-control activity (sidrophore, HCN and chitinase production) were statistically analyzed to determine if different growth forms of *D. strictus* have influenced the recovery of fluorescent pseudomonads isolates in a given category. Based on functional properties (growth and bio-control activities) of fluorescent pseudomonads, these properties were categorised into absent, weak, moderate and high and quantified as 1 (absent), 2 (weak), 3 (moderate) and 4 (high), respectively. The quantified data were further used for non-parametric tests, such as Kruskal–Wallis (using SPSS 19) to estimate the association between the various properties and growth forms of *D. strictus.* Pi-Chramer’s values were also calculated to estimate the size of association. The regression model was generated using colony and halozone diameters and as well as phosphate solubilising efficiency (XLSTAT, 2012). Multiple correspondence analyses were performed using XLSTAT 2012 to see the association between these microbial activities and geographical locations.

## 3. Results

### 3.1. Mycorrhizal Spore Population and Root Colonisation in different growth forms of *D. strictus*

The study registered seasonal variations in spore population of indigenous mycorrhizal population, for example, soil sample from the Shivpuri location had highest average spores (10.2/ml of soil, range 5.0–16.0) followed by FRI campus (9.3/ml of soil, range 5.0–13.7) and Chiriapur (8.7/ml of soil, range 3.7–12.7). There was a spike in spore numbers during the months of April and October and drop during the active monsoon month of July (Table 2; Figure 1A.). Some of the indigenous mycorrhizal spores from *D. strictus* soils are shown in Figure 2A and B.

**Table 2.** Spore population (no./ml of soil) of indigenous mycorrhizae in the root zone of different growth forms of *D. strictus*

| Site | Statistical parameter | Year I |  |  |  | Year II |  |  |  | Grand mean |
| --- | --- | --- | --- | --- | --- | --- | --- | --- | --- | --- |
|  |  | April | July | October | Mean | April | July | October | Mean |  |
| FRI Campus | Mean | 11.2 | 6.1 | 11.1 | 9.5 | 10.2 | 5.8 | 10.9 | 9.0 | 9.3 |
|  | Range | 8.3–13.7 | 5.0–8.0 | 8.3–13.7 |  | 7.7–11.3 | 5.0–7.0 | 7.7–13.3 |  |  |
| Shivpuri | Mean | 13.2 | 6.5 | 12.1 | 10.6 | 12.1 | 5.8 | 11.6 | 9.8 | 10.2 |
|  | Range | 10.7–16.0 | 5.3–7.7 | 9.0–15.0 |  | 10.0–14.0 | 5.0–6.7 | 9.7–12.3 |  |  |
| Chiriapur | Mean | 11.1 | 6.1 | 10.2 | 9.1 | 10.1 | 5.2 | 9.5 | 8.3 | 8.7 |
|  | Range | 10.0–12.7 | 4.7–7.3 | 8.7–12.0 |  | 8.7–11.0 | 3.7–7.0 | 8.0–10.3 |  |  |
|  | Mean | 11.8 | 6.2 | 11.1 | 9.7 | 10.8 | 5.6 | 10.7 | 9.0 | 9.4 |

**Figure. 1.**
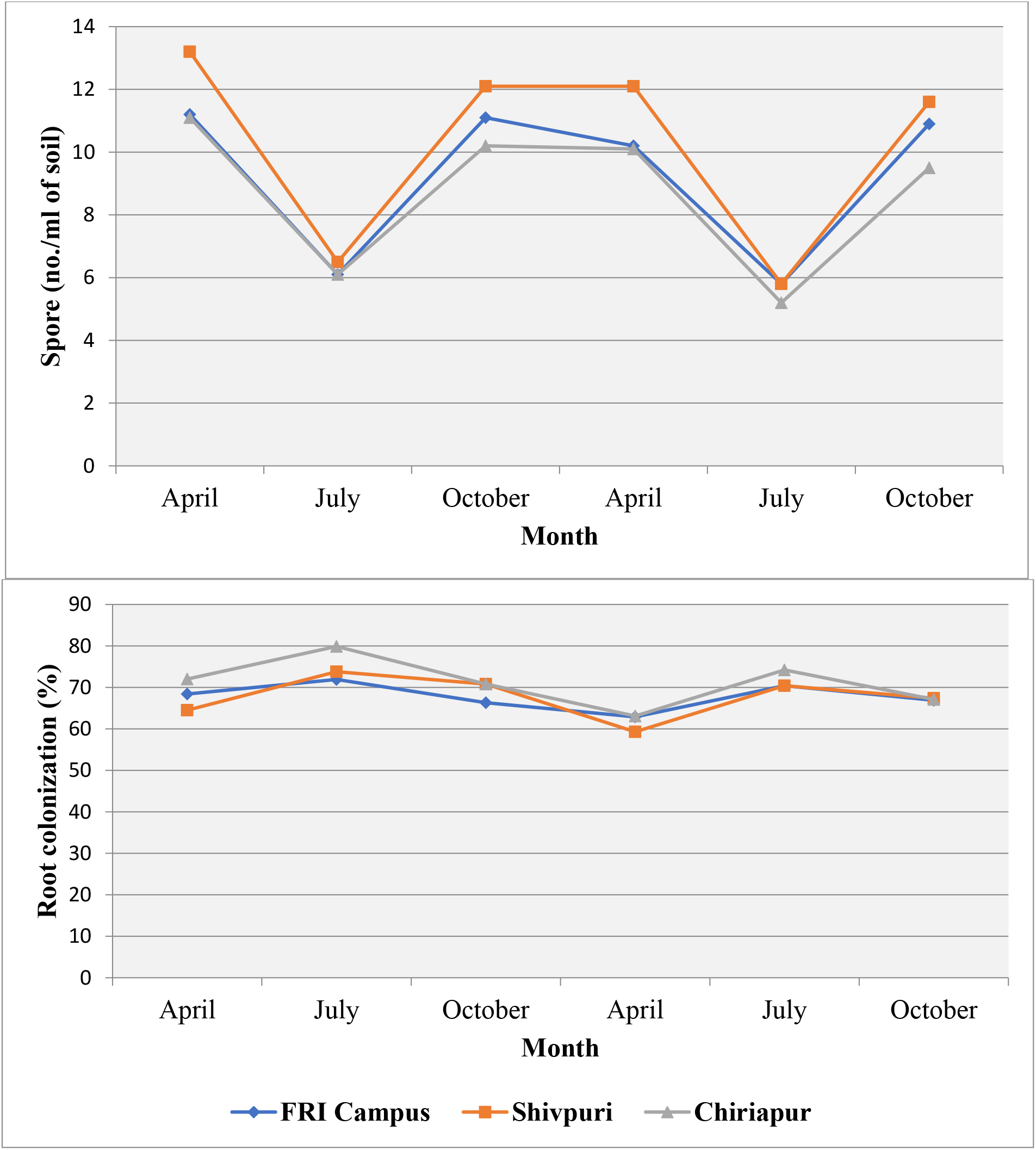
Seasonal changes in spore population (A) and root colonisation (B) of mycorrhizal fungi of different growth forms of *D. strictus*.

**Figure. 2.**
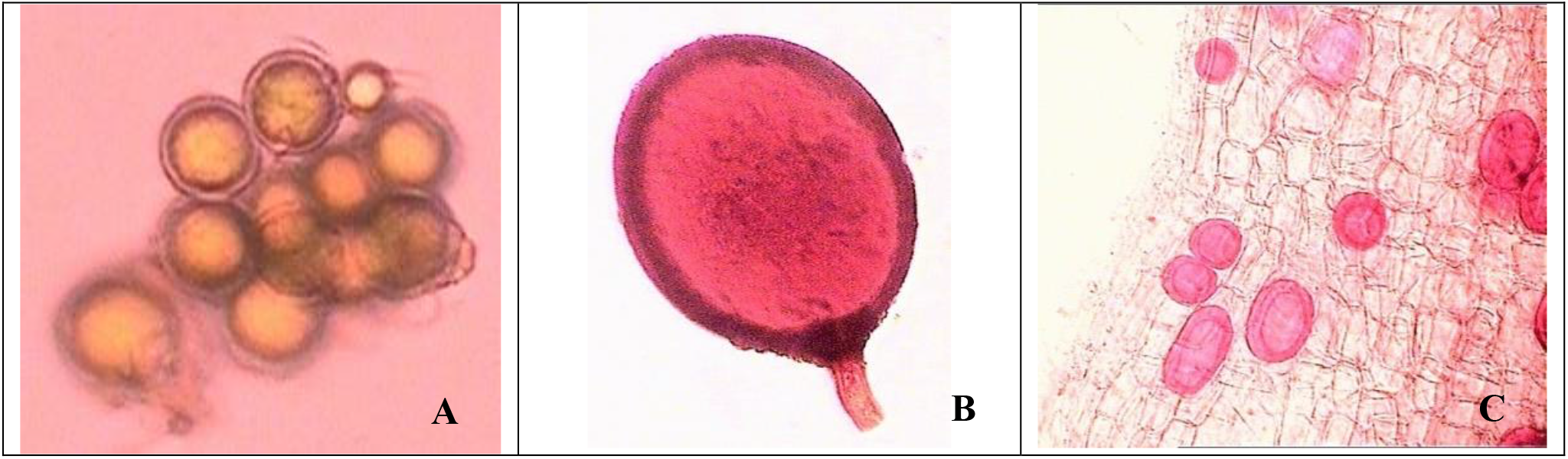
The indigenous endomycorrhizal spores (A and B) from soils and root colonisation by endomycorrhiza (vesicles, C) of different growth forms of *D. strictus*.

The root of *D. strictus* from Chiriapur site recorded the highest colonisation (71.2%, range 59.7– 92.8%), while FRI campus (67.8%, range 57.7–76.1%) and Shivpuri site (67.7%, range 54.7–82.3%) were close to each other (Table 3 and Figure 1B.). The root supporting some of the mycorrhizal colonizatons are shown Figure 2C.

**Table 3.**
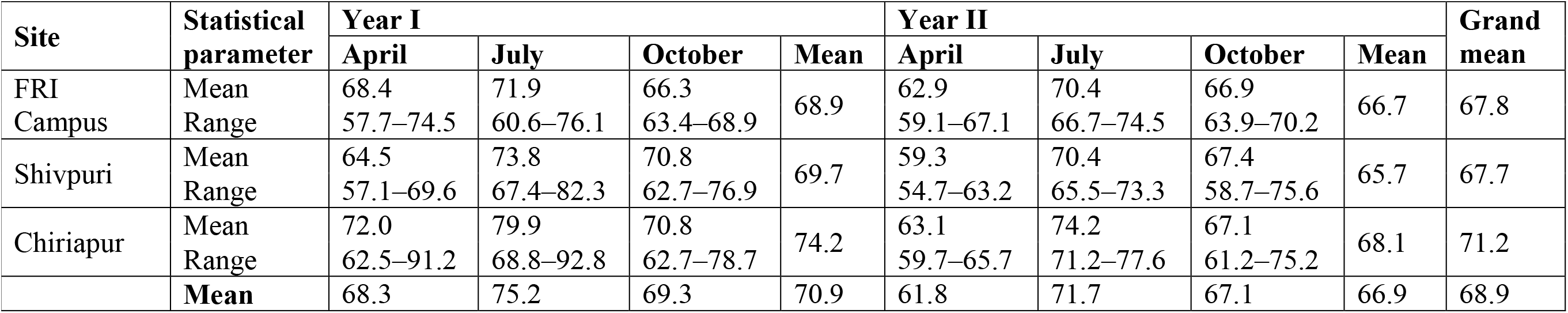
Root colonization (%) of *D. strictus* by indigenous mycorrhizae of different growth forms of *D. strictus*

| Site | Statistical parameter | Year I |  |  |  | Year II |  |  |  | Grand mean |
| --- | --- | --- | --- | --- | --- | --- | --- | --- | --- | --- |
|  |  | April | July | October | Mean | April | July | October | Mean |  |
| FRI Campus | Mean | 68.4 | 71.9 | 66.3 | 68.9 | 62.9 | 70.4 | 66.9 | 66.7 | 67.8 |
|  | Range | 57.7–74.5 | 60.6–76.1 | 63.4–68.9 |  | 59.1–67.1 | 66.7–74.5 | 63.9–70.2 |  |  |
| Shivpuri | Mean | 64.5 | 73.8 | 70.8 | 69.7 | 59.3 | 70.4 | 67.4 | 65.7 | 67.7 |
|  | Range | 57.1–69.6 | 67.4–82.3 | 62.7–76.9 |  | 54.7–63.2 | 65.5–73.3 | 58.7–75.6 |  |  |
| Chiriapur | Mean | 72.0 | 79.9 | 70.8 | 74.2 | 63.1 | 74.2 | 67.1 | 68.1 | 71.2 |
|  | Range | 62.5–91.2 | 68.8–92.8 | 62.7–78.7 |  | 59.7–65.7 | 71.2–77.6 | 61.2–75.2 |  |  |
|  | Mean | 68.3 | 75.2 | 69.3 | 70.9 | 61.8 | 71.7 | 67.1 | 66.9 | 68.9 |

### 3.2. Population Status and Functional Assessment of Fluorescent Pseudomonads from different*D. strictus* Growth Forms

A total of 343 isolates were collected from rhizosphere of three growth forms of *D. strictus* naturally occurring in different geographical sites in two years, i.e., Chiriapur site (127) followed by Shivpuri (119) and minimum from FRI campus (97; Table 4). The functional properties (growth – phosphate solubilising and production of IAA; biocontrol – siderophore, HCN and Chitinase production) of each isolate were categorised into four qualitative groups, namely, absent, weak, moderate and high. These categories were further used in resolving the functionality of isolates from three geographical sites.

**Table 4.**
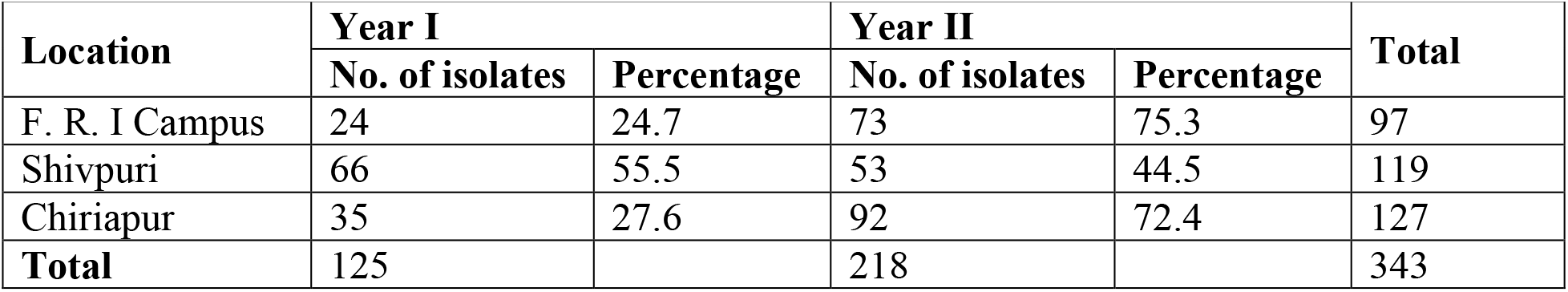
Population status of fluorescent pseudomonads isolated from different growth forms of *D. strictus*

| Location | Year I |  | Year II |  | Total |
| --- | --- | --- | --- | --- | --- |
|  | No. of isolates | Percentage | No. of isolates | Percentage |  |
| F. R. I Campus | 24 | 24.7 | 73 | 75.3 | 97 |
| Shivpuri | 66 | 55.5 | 53 | 44.5 | 119 |
| Chiriapur | 35 | 27.6 | 92 | 72.4 | 127 |
| **Total** | 125 |  | 218 |  | 343 |

#### 3.2.1. Phosphate Solubilising Activity

It was recorded that isolates from FRI campus site were poor phosphate solubilisers (79.4%) contrary to Chiriapur that contributed relatively high P solubilisers (11.8%; Table 5). Moreover, majority of the Shivpuri isolates were moderate solubilisers (84.0%).

**Table 5.** Status of plant growth promoting fluorescent pseudomonad isolates in the root zone of different growth forms of *D. strictus*

| Site | PS |  |  |  | IAA |  |  |  |
| --- | --- | --- | --- | --- | --- | --- | --- | --- |
|  | ABS | W | MOD | H | ABS | W | MOD | H |
| FRI Campus | 5 (5.2)* | 77 (79.4) | 14 (14.4) | 1 (1.0) | 21 (21.6) | 37 (38.1) | 29 (29.9) | 10 (10.3) |
| Shivpuri | 0 (0.0) | 19 (16.0) | 100 (84.0) | 0 (0.0) | 68 (57.1) | 28 (23.5) | 10 (8.4) | 13 (10.9) |
| Chiriapur | 4 (3.1) | 64 (50.4) | 44 (34.6) | 15 (11.8) | 61 (48.0) | 30 (23.6) | 27 (21.3) | 9 (7.1) |

\*Percentage values are in parentheses; PS-Phosphate Solubilising and IAA-Indole-3-Acetic Acid; ABS-Absent; W-Weak; MOD-Moderate; and H-High.

#### 3.2.2. Production of IAA

It was observed that majority of the isolates from Shivpuri site had no IAA activity (57.1%; Table 5) followed by Chiriapur isolates (48%). Furthermore, isolates of FRI campus had more moderate IAA activity (29.9%) as compared to others. Overall, 43.7% (Table 7) of 343 isolates showed no IAA activity, followed by weak (27.7%) and only 9.3% high IAA producers.

#### 3.2.3. Siderophore Production

The isolates from Shivpuri site had maximum poor siderophore producers (37.0%; Table 6), while the isolates from Chiriapur are moderate siderophore producers (40.2%). More than one-third isolates of FRI campus showed the highest siderophore activity (36.1%) as compared with others. Without taking sampling sites into consideration, overall, 36.4% isolates showed moderate, 29.2% weak and 27.4% high siderophore activity (Table 7).

**Table 6.**
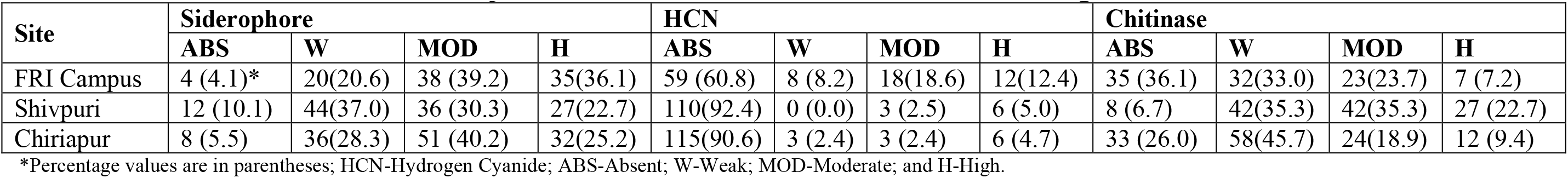
Status of biocontrol fluorescent pseudomonad isolates in the root zone of different growth forms of *D. strictus*

**Table 7.** Overall distribution of plant growth promoting and biocontrol activities of fluorescent pseudomonad isolates of different growth forms of *D. strictus*

| Category | PS | IAA | Siderophore | HCN | Chitinase |
| --- | --- | --- | --- | --- | --- |
| Absent | 9 (2.6)* | 150 (43.7) | 24 (7.0) | 284 (82.8) | 76 (22.2) |
| Weak | 160 (46.6) | 95 (27.7) | 100 (29.2) | 11 (3.2) | 132 (38.5) |
| Moderate | 158 (46.1) | 66 (19.2) | 125 (36.4) | 24 (7.0) | 89 (25.9) |
| High | 16 (4.7) | 32 (9.3) | 94 (27.4) | 24 (7.0) | 46 (13.4) |
| **Std. dev.** | ±0.59 | ±1.00 | ±0.91 | ±0.89 | ±0.91 |

\*Percentage values are in parentheses.

#### 3.2.3. HCN Activity

Irrespective of locations, majority of the bacterial isolates were HCN negative (82.8%). Out of which, isolates from Shivpuri showed no HCN activity contributing 92.4%, Chiriapur site 90.6% and FRI campus 60.8% (Table 6). However, it was interesting to note that about 12.4% of isolates from FRI site showed the highest HCN activity (Table 7).

#### 3.2.4. Chitinase Activity

A good number of Chiriapur isolates were weak chitinase producers (45.7%; Table 6) followed by Shivpuri (35.3%) and FRI campus (33%), while isolates from Shivpuri were good Chitinase producers (22.7%) as compared with FRI campus (7.2%) and Chiriapur (9.4%). Overall, 38.5% isolates showed weak chitinase activity (Table 7).

### 3.3 Kruskall–Wallis Test

The test was conducted on three populations of fluorescent pseudomonads. It was observed that except for chitinase activity (χ^2^ 4.37; df=2; *p*=0.112), correlation of all other characteristics (PSB, IAA, sidrophore and HCN) of these isolates were found to be mutually significant (Table 8). Based on the mean rank, the phosphate solubilising activity was weighed more in favour of Chiriapur (rank=212.5), while other properties like IAA (rank=207.0), siderophore (rank=196.0) and HCN (rank=208.7) production were found to be tilted more towards FRI campus.

**Table 8.** Mean ranking of different growth promoting and bio-control characteristics of fluorescent pseudomonad isolates of different growth forms of *D. strictus*

| Location | Characteristic | df<sup>@</sup> | N<sup>#</sup> | Mean rank | Chi-sq<sup>*</sup> | *P-value* |
| --- | --- | --- | --- | --- | --- | --- |
| FRI campus | PSB | 2 | 97 | 143.0 | 40.6 | 0.000 |
| Shivpuri |  |  | 119 | 152.4 |  |  |
| Chiriapur |  |  | 127 | 212.5 |  |  |
| FRI campus | IAA | 2 | 97 | 207.0 | 20.11 | 0.000 |
| Shivpuri |  |  | 119 | 151.6 |  |  |
| Chiriapur |  |  | 127 | 164.2 |  |  |
| FRI campus | SID | 2 | 97 | 196.0 | 11.4 | 0.003 |
| Shivpuri |  |  | 119 | 152.4 |  |  |
| Chiriapur |  |  | 127 | 172.0 |  |  |
| FRI campus | HCN | 2 | 97 | 208.7 | 42.93 | 0.000 |
| Shivpuri |  |  | 119 | 156.2 |  |  |
| Chiriapur |  |  | 127 | 158.9 |  |  |
| FRI campus | Chitinase | 2 | 97 | 159.8 | 4.37 | 0.112 |
| Shivpuri |  |  | 119 | 185.8 |  |  |
| Chiriapur |  |  | 127 | 168.5 |  |  |

<sup>@</sup>df = Degree of freedom; <sup>#</sup>N = Total number; <sup>\*</sup>Chi-sq value for df (2) = 5.9 at *P* 0.05 level of significance.

### 3.3. Multiple Correspondence Analysis (MCA)

Multiple correspondence analysis was conducted in order to see the association between different growth forms of *D. strictus* and functional properties (growth and bio-control) of fluorescent pseudomonads isolates. Overall, 58.64% variability was explained using two axes (F1 41.92%; F2 16.71%; Figure 3) of symmetric variable plot of MCA. It seems that Shivpuri (−0.895) and Chiriapur (0.894) laid on the same axis (F2) but in two opposite direction and FRI campus lying on F1 axis (1.211), which represented the variability among the growth forms. When growth and bio-control activities were considered in relation to pseudomonads populations, it was observed that high phosphate solubilising and chitinase activity were associated with isolates of Chiriapur site; the absence of siderophore and HCN activities were found to be associated with Shivpuri isolates; whereas, most of the high siderophore and HCN activities (bio-control) and high IAA activity were found to be associated with FRI campus isolates.

**Figure. 3.**
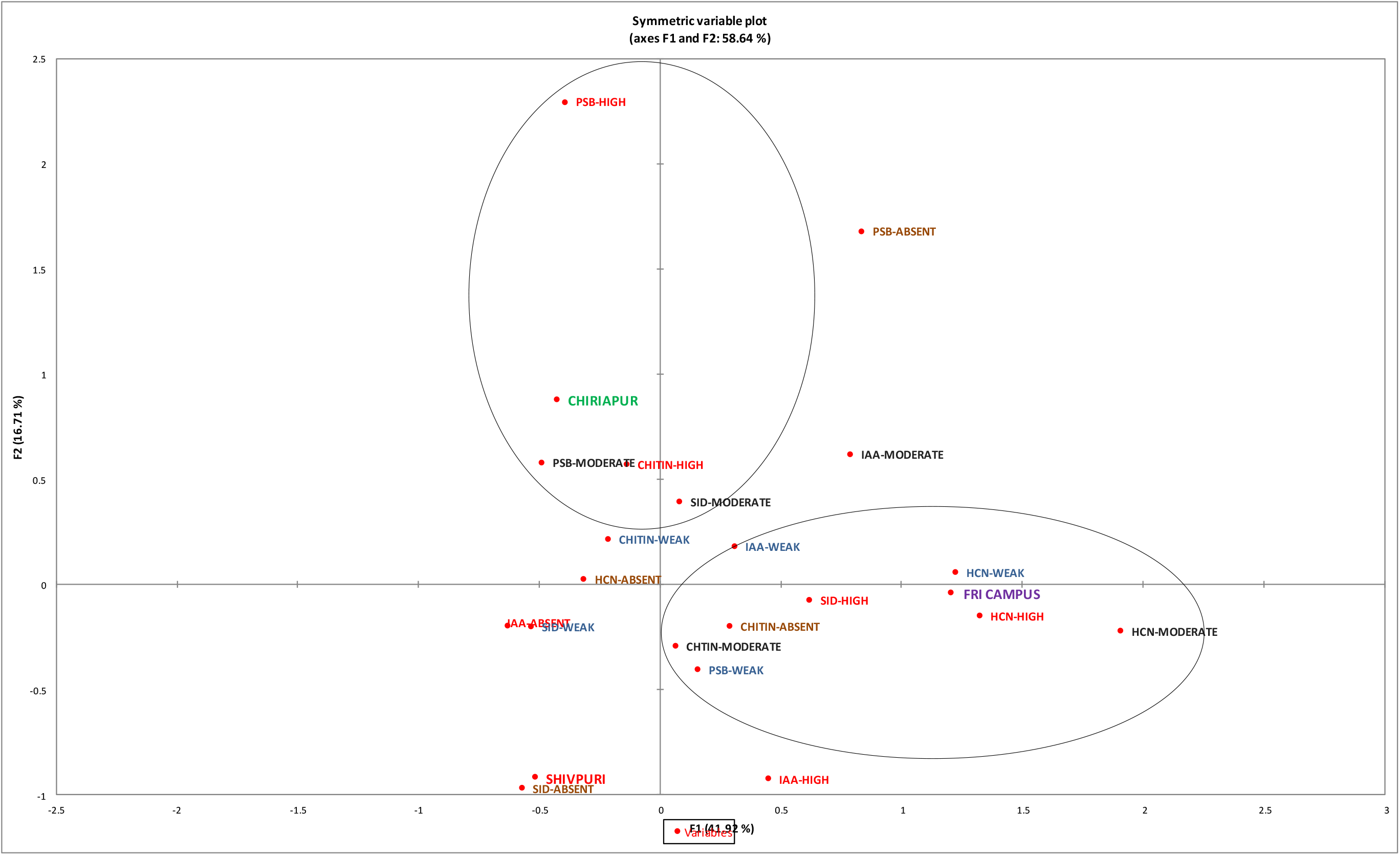
Multiple correspondence analysis of functional attributes of fluorescent pseudomonad isolates from the root zone of different growth forms of *D. strictus*.

### 3.4. Regression Model

Phosphate solubilising efficiencies (PSE) of isolates were estimated for each location and regression model was constructed using colony and halozone diameter as independent and PSE as dependent variable.

#### 3.4.1. FRI campus

The PSE was closely correlated with halozone (Pearson coefficient 0.740, R^2^=0.548) with 78.1% variability contributed by colony and halozone (R^2^=0.781). Hence, with the change in halozone diameter, the phosphate solubilising efficiency of the isolates will also be changed. The model was found to be statistically significant (F 168.0; *p*<0.0001) based on following equation:

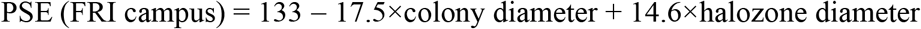

#### 3.4.2. Shivpuri

The PSE was correlated with halozone (Pearson coefficient 0.609, R^2^=0.370) with 96.5% variability contributed by colony and halozone (R^2^=0.964) in isolates of Shivpuri. The model was found to be statistically significant (F 1577.5; *p*<0.0001) with equation:

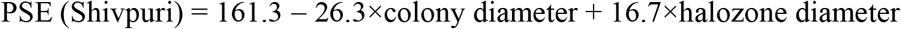

#### 3.4.3. Chiriapur

The PSE of Chiriapur was also closely correlated with halozone (Pearson coefficient 0.838, R^2^=0.702) around the colony with 95.4% variability contributed by colony and halozone (R^2^=0.954). The model was found to be statistically significant (F 1292.3; *p*<0.0001) with following equation:

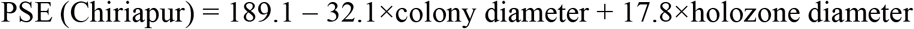

When all 343 isolates were clubbed together ignoring locations, it was found that PSE was closely related to halozone (Pearson coefficient 0.802) with 89.5% variability contributed by colony and halozone (R^2^=0.895). The model was statistically significant (F 1451.4; *p*<0.0001) with an equation:

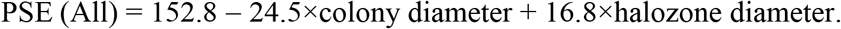

## 4. Discussion

### 4.1. Seasonal Survey of Bamboo Mycorrhiza and Population Status of Fluorescent Pseudomonads from *D. strictus* Growth Forms

*D. strictus*, being one of the prominent bamboo species in the country, plays a major role in supporting the socio-economic status of rural India. The different growth forms of *D. strictus* showed the morphological differences which may also find expression in the physiology of giant grass. Root-microbial interactions (mycorrhizzal as well as PGPRs) are crucial in the overall physiology of the host. The fast-growing nature and shallow root system of *D. strictus* create a gap between demand and supply of the nutrients especially P in tropics that may be filled with the use of beneficial microbes, mainly solubiliser (fluorescent pseudomonads) and absorber (AM fungi).

Seasonal survey reveals the microbial status (AM +fluorescent pseudomonads) in the rhizospheres of different growth forms of bamboo. It is important to understand the functionality and its extent as the bacterium is free living in the root zone of the bamboo while, the fungus affects the host physiology through direct colonization of the roots. The endomycorrhizal spore population and root colonization of different growth forms of *D. strictus* showed variation in different seasons (Tables 2 and 3). There was a drop in spore population during rainy season (July) from April (onset of summer) and then, boost with the onset of winter (October). Similar trends were observed for the consecutive years. The percent root colonization, however, increased during the rainy season and decreased with the onset of winter. These trends were similar to the observation of Rawat (2005), who studied the seasonal variations of endomycorrhizal spores and root colonization from the rhizospheres of same growth forms of *D. strictus*. The trend of spore population in present study was in contradiction with the findings of Sharma et al. (2005), who also studied the seasonal dynamics of arbuscular mycorrhizal fungi of *D. strictus* of Haryana. They reported that there was a steady increase in spore population from summer to rainy season with the maximum spores during the rainy season. In the present case, the maximum root colonization during rainy season might be due to high root proliferation and rhizome formation. Besides, creating a higher demand for nutrients vis-a-vis biomass gathering, it might also be providing more entry points for AM fungi (Bhaskaran and Selvaraj, 1997; Allen *et al.* 1998) leading to higher colonization. As root colonization is an outcome of spore germination and, subsequent, germ tube entering, fall in spore number during the monsoon is a natural consequence of higher colonization. Similarly, there was a decrease in spore population in *D. strictus* root zone with the onset of winter. It might be due to the beginning of senescence phase of the host when root growth and proliferation was at decrease. The physical manifestation of symbiosis in form of extensive root colonization of *D. strictus* by indigenous fungi present at different sites supported the early assumption that the host must be highly mycotrophic due to its shallow root system and fast rate of growth. Moreover, the relationship between myorrhizal association and growth pattern of *D. strictus* can also not be ignored. During spring season, a profuse leafing marked the growth initiation phase of this bamboo with accumulation of food during summers. These properties, later, led to the formation of new rhizomes and tillering in the monsoon. While in winter, shedding of leaf reflects the low metabolic activity of the host. All these physiological stages of *D. strictus* in different seasons were very much supported by the extent of indigenous mycorrhizal root colonization.

When the soil P and organic matter were taken into considerations to understand the role of these soil factors in the process of mycorrhization, the soil P (12.5 kg/ha) and organic matter (0.97%) of Chiriapur were on the lower side as compared to Shivpuri (53.8 kg/ha; 1.98%) and FRI campus (69.9 kg/ha; 2.4%; Table 1). These observations are in corroboration of the findings of Kobae et al. (2016) who noted that with the increase in P treatment, there was a significant reduction in arbuscule density as well as in mycorrhizal hyphae. They also suggested that there was a negative effect of P on AM root colonization. An increase in soil P is expected to bring about certain anatomical and physiological modifications in the plant roots which could restrict the spread of intra-radical AM fungi (Menge, *et al.*, 1978; Amijee *et al.*, 1989; Nagahashi *et al.*, 1996). Therefore, a good AM colonization in Chiriapur appeared to be a result of low soil P. Having maximum soil P and organic content in FRI campus, *D. strictus* at this location had the minimum root colonization for both the years. Douds (1994) also observed that high soil P content restricted the supply of organic carbon from roots to AM fungus, which, in return, reduced the proliferation of mycorrhizal hyphae and formation of spores.

Fluorescent pseudomonads population is present in large numbers in all natural environments, like terrestrial, freshwater and marine. They are known to have several important functions, of which plant growth promotion and soil nutrient cycling are of our interest. The presence of these bacteria along with mycorrhizae supported the concept of Garbaye (1994), who reported that fluorescent pseudomonads as one of the PGPRs (plant growth promoting rhizobacteria) that could also act as ‘mycorrhizal helper bacteria’. The contradictory trend in recovery of fluorescent pseudomonads populations between FRI campus and Chiriapur might be an outcome of sampling and isolation limitations.

### 4.2. Assessment of Functional Characteristics of Fluorescent Pseudomonads Population from *D. strictus* Rhizospheres

Plant rhizosphere is the seat of microbial diversity. These microbes are essential for different soil ecological processes, such as recycling of essential elements (carbon, phosphorus, nitrogen and sulphur), decomposition of organic matter, etc, and hence, directly or indirectly influence lives on earth (Garbeva *et al.*, 2004). Fluorescent pseudomonad populations isolated from rhizospheres of three growth forms of *D. strictus* were functionally characterised and investigated to find out whether functional variability among and within bacterial populations of different growth forms existed or not? *In-vitro* qualitative analysis of growth and biocontrol properties reflected the functional variability among and within the isolates of different locations. The present study revealed that except for cyanide production, majority of the indigenous isolates of fluorescent pseudomonads were involved in phosphate solubilisation, synthesis of phytohormone (IAA), siderophore production and chitinase activity (Tables 5, 6 and 7). The absence of HCN production in majority of the isolates indicated that rhizospheres of *D. strictus* had favoured non-HCN producers, which, in turn, are useful for unrestricted growth of shallow root system of bamboo. As explained by Kapulnik (1996), the cyanide produced by microorganisms is having inhibitory effect over ATP (respiratory energy) producing ability of roots. This may also reduce the efficiency of roots for nutrient uptake. Both these conditions go against the fast rate of growth of bamboo that can only be achieved by the plant when its, otherwise, shallow root system remains healthy and efficient in its functions including nutrient uptake. Therefore, the plant prefers such class of PGPR in its root zone that either do not have or have poor in root inhibition factors like HCN production.

Based on Kruskall-Wallis test (Table 8), it could be inferred that the efficient phosphate solubilisers were more common in Chiriapur, while isolates from FRI campus were better IAA, sidrophore and HCN producers. The MCA (Figure 3) placed the efficient (high) P solubilisers in the realm of Chiriapur prossibly due to P deficient soil. Moreover, population of fluorescent pseudomonads producing high siderophore and HCN were found to be associated with FRI campus where organic matter rich soil was prevalent boosting the soil microbial activity due to readily available food in the form of organic matter. This situation may lead to intense microbial competition for the food leading to development of various mechanism of self-defence (siderophore, HCN and chitinase) in the indigenous fluorescent pseudomonads populations. Soil fluorescent *Pseudomonas* species from undisturbed sites (forests/plantations) are not universally distributed (Cho and Tiedje 2000) indicating thereby a constant diversification and adaptation to local conditions. These observations are further supported by the facts that different soil conditions like, soil pH, organic matter and soil nutrients (NPK) may be influencing the composition and functional attributes of these bacteria (Kwon *et al.*, 2005). For example, sandy loamy soil and poor soil P of Chiriapur tended to support efficient P solubilisers. The indigenous population of bacteria from this location could have adapted to nutrient deficient condition. While fluorescent pseudomonads of FRI campus, where soils are nutrient (in terms of soil P and organic matter) and organisms (fungi and other bacteria) rich may have evolved to support more siderophore producers to overcome microbial competition. Such functional variations in fluorescent pseudomonads might also be due to the difference in composition of root exudates as well as different ecological zones.

Regression models are used to predict one dependent variable from one or more independent variables. A fitness of model was examined by coefficient of determination (R^2^) and model having R^2^-value higher than 0.9 indicated high predictability (Chen *et al.*, 2009). Except for FRI campus (R^2^ =0.78), both Shivpuri (R^2^ =0.96) and Chiriapur (R^2^ =0.95) represented a very good fit between the observed and predicted response. The higher β-coefficient value for Chiriapur (β=189.1) reflected that isolates from this location had higher halozone diameter, which directly indicated the presence of better P solubilisers. Eventually, the present study is directing towards the support of three phenomena, which Lemanceau *et al.* (1995) gave in relation to the effect of two plant species on the diversity of soil-borne populations of fluorescent pseudomonads. They envisaged that (1) plants play a selective influence on fluorescent pseudomonads populations, (2) the degree of selection depends upon the host plant or varies with respect to host plant, and (3) selection of bacterial population is partially plant specific. These three postulates can be validated by the present study. For example, the first one can be explained or supported based on the functional diversity of fluorescent pseudomonads from these growth forms. It was observed that the variations in functionality of these bacteria, like better P solubilisers from Chiriapur, high siderophore producers from FRI campus and absence of HCN producers from Shivpuri indicated that respective growth form had shaped the fluorescent pseudomonad populations in its rhizosphere. Second postulate is verified with one functional characteristic of the bacterium i.e., irrespective of growth forms, *D. strictus* did not support the population of fluorescent pseudomonads having HCN activity (in all 82.8% isolates) as it is directly beneficial for normal root growth of the plant that has, otherwise, shallow root system. Fast growth rate of bamboo required a healthy and functional root system. Thirdly, the nutrient poor soil in Chiriapur (especially P deficient) challenged both host as well as its rhizospheric pseudomonads population as both the organisms had P requirements for their normal growth. The selection pressure of nutrient deficient condition might have created a situation that could have given rise to the occurrence of high numbers of efficient P solubilisers which, in turn, also boosted the bamboo growth. This common challenge to both plant and its microbe primarily influenced the microbe and limits the role of host in the selection.

## Conclusion

The study revealed the complexity of dynamics played between bamboo root system and microbial web. The shallow root system of bamboo seems to be depending on the symbiotic associations of indigenous arbuscular mycorrhizae and fluorescent pseudomonas population for the uptake of nutrients for its faster growth. There seems to be influence of environment, soil ecology, growth forms of *D. strictus*, and soil P on the population of mycorrhizal population and functional diversities of fluorescent pseudomonas population. Though further study is required to establish more direct effect as well as to explore the much more complexities of plant-microbe interactions, especially in relation with the faster growing plants such as bamboo.

## Acknowledgement

We would like to thank all the research staff of Forest Pathology Division for their constructive support in this study. This study is not funded by any organisation.

## Author Contributions

Research design, concept, writing of manuscript, statistical analyses, and experiments were conducted by Solomon Das. Concept, experimental design, and editing by Y.P. Singh. Research protocol, correction, overall conceptualization by Yogesh Negi. Soil experiments, research design, and conceptualization by P. C. Shrivastava.

## References

Ahmad, F.; Ahmad, I. and Khan, M.S. 2005. Indole acetic acid production by the indigenous isolates of azotobacter and fluorescent in the presence and absence of tryptophan. Turk. J. Biol., 29:29–34.

Allen, E.B.; Rincon, E.; Allen, M.F.; Perez-Jimenez, A. and Huante, P. 1998. Distribution and seasonal dynamics of mycorrhizae in a tropical deciduous forest in Mexico. Biotropical., 30:261–274.

Amijee, F.; Tinker, P.B. and Stribley, D.P. 1989. The development of endomycorrhizal root systems. VII. A detailed study of effects of soil phosphorus on colonization. New Phytol, 111 (3): 435–446.

Bhaskaran, C. and Selvaraj, T. 1997. Season incidence and distribution of VA–mycorrhizzal fungi in native saline soils. J. Environ. Biol., 18:209–212.

Bric, J.M.; Bostock, R.M. and Silverstone, S.E. 1991. Rapid *in situ* assay for indole acetic acid production by bacteria immobilized on nitrocellulose membrane. Appl. Environ. Microbiol., 57:535–538.

Chakraborty, R N.; Patel, H.N. and Desai, S B. 1990. Isolation and partial characterization of catechol-type siderophore from *Pseudomonas stutzeri* RC-7. Curr. Microbiol., 20: 283–286.

Chen, X.C.; Bai, J.X.; Cao, J.M.; Li, Z.J.; Xiong, J.; Zhang, L.; Hong, Y. and Ying, H J. 2009. Medium optimization for the production of cyclic adenosine 3’, 5’-monophosphate by *Microbacterium* sp. no. 205 using response surface methodology. Bioresour. Technol., 100: 919–924.

Cho, J.C and Tiedje, J.M. 2000. Biogeography and degree of endemicity of fluorescent Pseudomonas strains in soil. Appl. Environ. Microbiol., 66 (12): 5448.

Daniels, B A. and Skipper, H.D. 1982. Methods for the recovery and quantitative estimation of propagules from soil. In: Schenck, N.C. (ed.). Methods and principles of mycorrhizal research. St. Paul, MN, The American Phytopathological Society. pp.29–36.

Das. S.; Singh, Y.P.; Negi, Y K. and Shrivastav, P.C. 2017. Genetic variability in different growth forms of *Dendrocalamus strictus*: Deogun revisited. New Zealand Journal of Forestry Science, 47: 23–34. .DOI 10.1186/s40490-017-0104-4

Deogun, P.N. 1937. The silviculture and management of the bamboo *Dendrocalamus strictus* Nees, 2 (4):173.

Didick H.G.; Siswanto and Sugiarto, Y. 2000. Bioactivation of poorly soluble phosphate rocks with a phosphorus solubilizing fungus. Soil Sci. Soc. Am. J., 64: 927–932.

Douds, D. D. 1994. Relationship between hyphal and arbuscular colonization and sporulation in a mycorrhiza of *Paspalum notatum* Flugge. New Phytol., 126 (2): 233–237.

Fankem, H.; Nwaga, D.; Deubel, A.; Dieng, L.; Merbach, W. and Etoa, F.X. 2006. Occurrence and functioning of phosphate solubilising microorganisms from oil palm tree rhizosphere in Cameroon. Afr. J. Biotechnol., 5 (24): 2450–2460.

Garbaye, J. 1994. Mycorrhiza helper bacteria: A new dimension to the mycorrhizal symbiosis. New Phytol., 128: 197–210.

Garbeva, P.; van Veen, J.A. and van Elsas, J.D. 2004 Assessment of the diversity, and antagonism towards *Rhizoctonia solani* AG3, of *Pseudomonas* species in soil from different agricultural regimes. FEMS Microbiol. Ecol., 47: 51–64.

Gerdemann, J.W. and Nicolson, T.H. 1963. Spores of mycorrhizal Endogone extracted from soil by wet sieving and decanting. Trans. Brit. Mycol. Soc., 46: 235–244.

Giovannetti, M. and Mosse, B. 1980. An evaluation of techniques for measuring vesicular-arbuscular mycorrhizzal infection in roots. New Phytol., 84: 489–500.

Gloria, W.A. and Hoadley, A.W. 1976. Fluorescent pseudomonads capable of growth at 41°C but distinct from *Pseudomonas aerugenosa*. J. Clin. Microbiol., 4: 443–449.

INBAR. 2018, A new report by INBAR, the Food and Agriculture Organization of the UN and the New Partnership for Africa’s Development looks at bamboo’s use to restore degraded lands in eight places across the world: China, Colombia, Ghana, India, Nepal, South Africa, Tanzania and Thailand, June 18, 2018.

Kapulnik, Y. 1996, Plant growth promoting rhizosphere bacteria, In: Waisel, Y.; Eshel, A. and Kafkafi, U. (eds.). Plant roots: The hidden half. New York, Marcel Dekker. pp.769–781.

Kobae, Y.; Ohmori, Y;, Saito, C.; Yano, K.; Ohtomo, R. and Fujiwara, T. 2016. Phosphate treatment strongly inhibits new arbuscule development but not the maintenance of arbuscule in mycorrhizal rice roots. Plant Physiology, 171(1): 566–579.

Kwon, S.W.; Kim, J.S.; Crowley, D.E. and Lim, C.K. 2005. Phylogenetic diversity of fluorescent pseudomonads in agricultural soils from Korea. Lett. Appl. Microbiol., 41: 417–423.

Lemanceau, P.; Corberand, T.; Gardan, L.; Latour, X.; Laguerre, G.; Boeufgras, J.M. and Alabouvette, C. 1995. Effect of two plant species, flax (*Linum usitatissinum* L.) and tomato (*Lycopersicon esculentum* Mill.) on the diversity of soil-borne populations of fluorescent pseudomonads. Appl. Environ. Microbiol., 61:1004–1012.

Menge, J.A.; Sterile, D.; Bagyaraj, D.J.; Johnson, E. L.V. and Leonard, R.T. 1978. Phosphorus concentrations in plants responsible for inhibition of mycorrhizal infection. New Phytol., 80 (3):575–578.

Meyer, J.M. and Abdallah, M.A. 1978. The fluorescent pigment of *Pseudomonas fluorescens* biosynthesis, purification and physical-chemical properties. J. Gen. Microbiol., 107:319–328.

Millar, R.L and Higgins, V. J. 1970. Association of cyanide with infection of birds foot trefoil by *Stemphylium loti*. Phytopathology, 60: 104–110.

Nacoon, S.; Jogloy, S.; Riddech, N.; Mongkolthanaruk, W.; Kuyper, T.W. and Boonlue, S. 2020. Interaction between phosphate solubilizing bacteria and arbuscular mycorrhizal fungi on growth promotion and tuber inulin content of *Helianthus tuberosus* L. Scientific Reports, 10: 4916.

Nagahashi, G.; Douds Jr., D. D. and Abney, G.D. 1996. Phosphorus amendment inhibits hyphal branching of the VAM fungus, *Gigaspora margarita* directly and indirectly through its effect on root exudation. Mycorrhiza, 6 (5): 403–408.

NMBTTD, 2003. Report. New Delhi, Planning Commission.

Phillips, J.M. and Hayman, D.S. 1970. Improved procedures for clearing roots and staining parasitic and vesicular-arbuscular mycorrhizal fungi for rapid assessment of infection. Trans. Brit. Mycol. Soc., 55: 158–161.

Pikosvskaya, R. I. 1948. Mobilization of phosphorus in soil in connection with vital activity of some microbial species. Microbiology, 17: 362–370.

Rawat, S. 2005. Bamboo-endomycorrhiza: Ecology, growth and macroproliferation. Ph. D. thesis, Forest Research Institute (Deemed) University, Dehradun. 200p.

Renwick, A.; Campbell, R. and Coe, S. 1991. Assessment of *in vivo* screening systems for potential biocontrol agents of *Gaeumannomyces graminis*. Plant Pathol., 40: 524–532.

Rijavec, T., and Lapanje, A. 2016. Hydrogen Cyanide in the rhizosphere: Not suppressing plant pathogens, but rather regulating availability of phosphate. Frontiers in Microbiology, 7: 1785.

Santana, E.B., Marques, E.L.S., and Dias, J.C.T., 2016. Effects of phosphate-solubilizing bacteria, native microorganisms and rock dust on *Jatropha curcas* L. growth. Genetics and Molecular Research, 15 (4): gmr.15048729.

Schenck, N.C. and Perez, Y. 1990. Manual for the identification of VA mycorrhizal fungi. Gainesville, Fl, Synergistic Publications. 123p.

Sharma, S.; Aggarwal, A.; Parkash, V. and Mehrotra, R.S. 2005. Seasonal population dynamics of vesicular arbuscular mycorrhizal fungi associated with *Tectona grandis* L. and *Dendrocalamus strictus* (Roxb.) Nees. Indian Phytopathol., 58 (2): 163–166.

Thangavelu, M., and Udaiyan, K. 2006. Growth of nursery-grown bamboo inoculated with arbuscular mycorrhizal fungi and plant growth promoting rhizobacteria in two tropical soil types with and without fertilizer application. New Forests, 31(3): 469–485.

